# Understanding sequence conservation with deep learning

**DOI:** 10.1101/103929

**Authors:** Yi Li, Daniel Quang, Xiaohui Xie

## Abstract

**Motivation:** Comparing the human genome to the genomes of closely related mammalian species has been a powerful tool for discovering functional elements in the human genome. Millions of conserved elements have been discovered. However, understanding the functional roles of these elements still remain a challenge, especially in noncoding regions. In particular, it is still unclear why these elements are evolutionarily conserved and what kind of functional elements are encoded within these sequences.

**Results:** We present a deep learning framework, called DeepCons, to uncover potential functional elements within conserved sequences. DeepCons is a convolutional neural net (CNN) that receives a short segment of DNA sequence as input and outputs the probability of the sequence of being evolutionary conserved. DeepCons utilizes hundreds of convolution kernels to detect features within DNA sequences, and automatically learns these kernels after training the CNN model using 887,577 conserved elements and a similar number of nonconserved elements in the human genome. On a balanced test dataset, DeepCons can achieve an accuracy of 75% in determining whether a sequence element is conserved or not, and the area under the ROC curve of 0.83, based on information from the human genome alone. We further investigate the properties of the learned kernels. Some kernels are directly related to well-known regulatory motifs corresponding to transcription factors. Many kernels show positional biases relative to transcriptional start sites or transcription end sites. But most of discovered kernels do not correspond to any known functional element, suggesting that they might represent unknown categories of functional elements. We also utilize DeepCons to annotate how changes at each individual nucleotide might impact the conservation properties of the surrounding sequences.

**Availability:** The source code of DeepCons and all the learned convolution kernels in motif format is publicly available online at https://github.com/uci-cbcl/DeepCons.

## 1 Introduction

One of the most surprising discovery of sequencing and comparing the human and other mammalian genomes is the revelation of conserved sequences, most of which are located outside of genes. Some of these conserved sequences exhibit few nucleotide changes throughout millions of years mammalian evolution, suggesting strong purifying selection within these sequences [1, 2]. Studies based on human and rodent genomes estimate that about 5% bases of mammalian genomes are under negative selection, among which coding regions only account for 1.5% of the genome [1]. Extensive studies and methods have been focused on understanding the functional roles of conserved sequences in noncoding regions. Nevertheless, the exact function of conserved non-coding sequences remains elusive.

Recent advances in deep learning, specifically in solving sequence-based problems in genomics with convolutional neural networks [3, 4, 5, 6], provide a new powerful method to study sequence conservation. Deep learning refers to algorithms that learn a hierarchical nonlinear representation of large datasets through multiple layers of abstraction (e.g. convolutional neural networks, multi-layer feedforward neural networks, and recurrent neural networks). It has achieved state-of-the-art performances on several machine learning applications such as speech recognition [7], natural language processing [8], and computer vision [9]. More recently, deep learning methods have also been adapted to solve genomics problems such as motif discovery [4, 5, 6], pathogenic variants identification [10], and gene expression inference [11].

In this paper we present a deep learning method for studying sequence conservation (DeepCons). DeepCons is a convolutional neural network trained to predict whether a given DNA sequence is conserved or not. By learning to discriminate between conserved and non-conserved sequences, DeepCons can capture rich information about conserved sequences, such as motifs. Specifically, we show that, 1) some of the learned convolution kernels are directly related to known motifs, such as regulatory motifs CTCF and the RFX family, that are known to be widely distributed within conserved noncoding elements [12], 2) lots of the kernels have positional bias relative to transcription start sites (TSS), transcription end sites (TES) and miRNA, indicating their potential roles in transcriptional control and post-transcriptional regulation, and 3) the kernels that are close to TES display strand bias, suggesting their RNA level regulatory effects. We further demonstrate that DeepCons could be used to score sequences at nucleotide level resolution in terms of conservation. We rediscovered known motifs, such as CTCF, JUND, RFX3 and MEF2A, within a given sequence by highlighting each nucleotide regarding their scores. Finally, we show that the learned convolution kernels represents a large variety of motifs, and we have made all the kernels publicly available online in the MEME [13] format. We hope researchers may draw new biological insights from these motifs.

## 2 Methods

### 2.1 DeepCons

DeepCons is a convolutional neural network [3] composed of one input layer, three hidden layers and one output layer. The first input layer uses one hot encoding to represent each input sequence as a 4-row binary matrix, with the number of columns equal to the length of the sequence. The second layer is a convolution layer composed of different convolution kernels with rectified linear units as the activation function. Each convolution kernel acts as a motif detector that scans across input matrices and produces different strengths of signals that are correlated to the underlying sequence patterns. The third layer is a max pooling layer that takes the maximum output signals of each convolution kernel along the whole sequence. The fourth layer is a fully connected layer with rectified linear units as activation. The last layer performs a non-linear transformation with sigmoid activation and produces a value between 0 and 1 that represents the probability of a sequence being conserved. DeepCons contains ∼2.3 million parameters. Figure 1 shows the neural network architecture of DeepCons.

**Figure 1:**
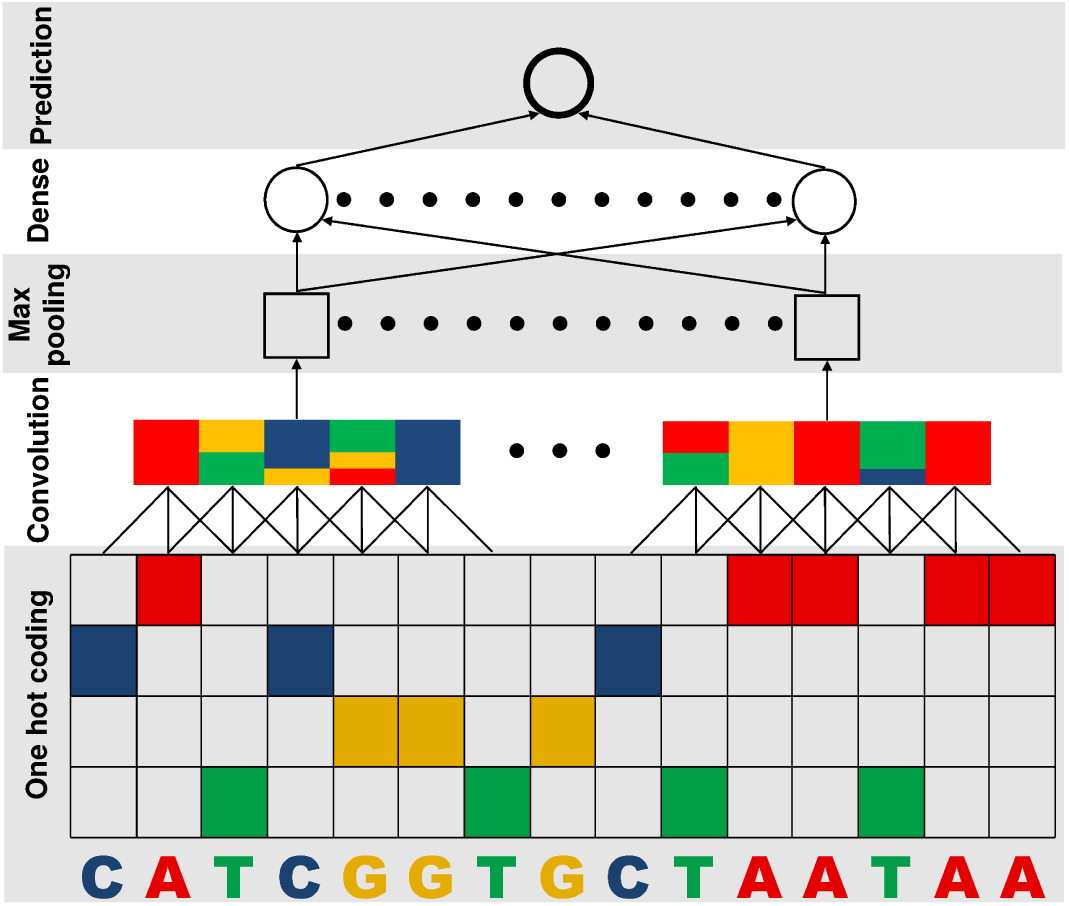
The neural network architecture of DeepCons.

DeepCons was trained using the standard back-propagation algorithm [14] and mini-batch gradient descent with the Adagrad [15] variation. Dropout [16] and early stopping were used for regularization and model selection. Detailed parameter configurations are given in the Supplementary.

DeepCons was implemented based on two Python libraries, Theano [17] and Keras http://keras.io/. Training was performed on an Nvidia GTX TITAN Z graphics card. DeepCons is publicly available at https://github.com/uci-cbcl/DeepCons.

### 2.2 Logistic regression

We also trained a baseline model using logistic regression (LR) for benchmarking purposes. Instead of using raw sequences as inputs, we first computed the counts of different k-mers of length 1-5 bp [4]. We then normalized the counts by subtracting the mean and dividing by the standard deviation and used these values as features. We also added a small L2 regularization of 1*e*-6 to the cross entropy loss function of LR during training. LR was implemented with the scikit-learn [18] library.

## 3 Results

In this section, we first introduce the dataset of conserved and non-conserved sequences we used in this study and show the performances of both DeepCons and LR on this dataset. Next, we demonstrate that DeepCons captures rich information within the conserved sequences by showing that some learned convolution kernels correspond to known motifs, many kernels have positional bias relative to TSS, TES and miRNA, and display strand bias relative to TES. We further demonstrate that the learned model could be used to score the importance of each nucleotide within a given sequence in terms of conservation, and rediscovered known motifs with these scores. Finally, we clustered all the convolution kernels and show that they represents a large variety of informative motifs.

### 3.1 Classifying conserved and non-conserved sequences

We first show that DeepCons can accurately discriminate between conserved and non-conserved sequences. To build the dataset of conserved sequences, we downloaded the 46-way phastCons conserved elements [1] under mammal category from UCSC genome browser [19] based on hg19. We excluded conserved sequences that overlap with repetitive sequences (http://www.repeatmasker.org/). or coding exons. We then filtered away conserved sequences that were either shorter than 30 bp or longer than 1,000 bp for training the model, leaving 887,577 sequences in the end. 75% of the nucleotides were preserved after the length filtering. To build the dataset of non-conserved sequences, we randomly shuffled the 887,577 conserved sequences on hg19, excluding repetitive sequences, coding exons and conserved sequences themselves. After combining both conserved and non-conserved sequences, we randomly set aside ∼80% for training (1,415,154 sequences), ∼10% for validation (180,000 sequences) and ∼10% for testing (180,000 sequences). Details of data preprocessing are given in the Supplementary.

DeepCons achieved 74.9% accuracy and an area under the curve (AUC) of 0.830 on the testing dataset. The baseline LR model achieved 65.9% accuracy and 0.722 AUC on the testing dataset. Figure 2 shows the receiver operating characteristic (ROC) curves of both DeepCons and LR on the testing dataset. DeepCons outperforms the baseline LR model significantly on classifying conserved and non-conserved sequences.

**Figure 2:**
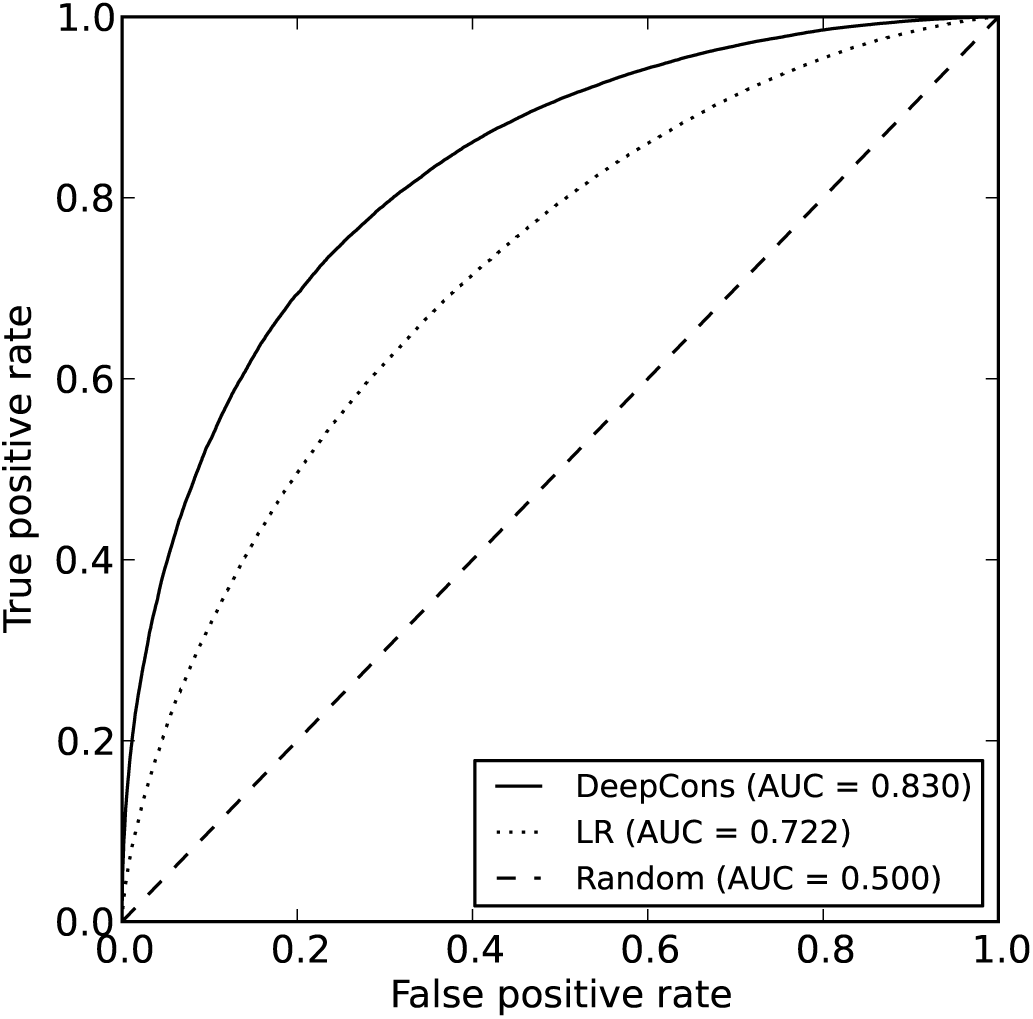
The ROC curves of DeepCons and LR on classifying conserved and non-conserved sequences on the testing dataset.

#### 3.2 Known motifs

Previous results have shown that regulatory motifs are widely distributed within conserved noncoding elements across the human genome, such as CTCF and the RFX family [12]. We observed similar results when examining the learned convolution kernels of DeepCons. Specifically, we converted the kernels from the convolution layer to position weight matrices, using the method described in DeepBind [5]. Then, we aligned these kernels to known motifs using TOMTOM [20]. 69 kernels match known motifs significantly (*E* < 1*e* – 2), including the CTCF and RFX families. Figure 3 shows four examples of identified known motifs.

**Figure 3:**
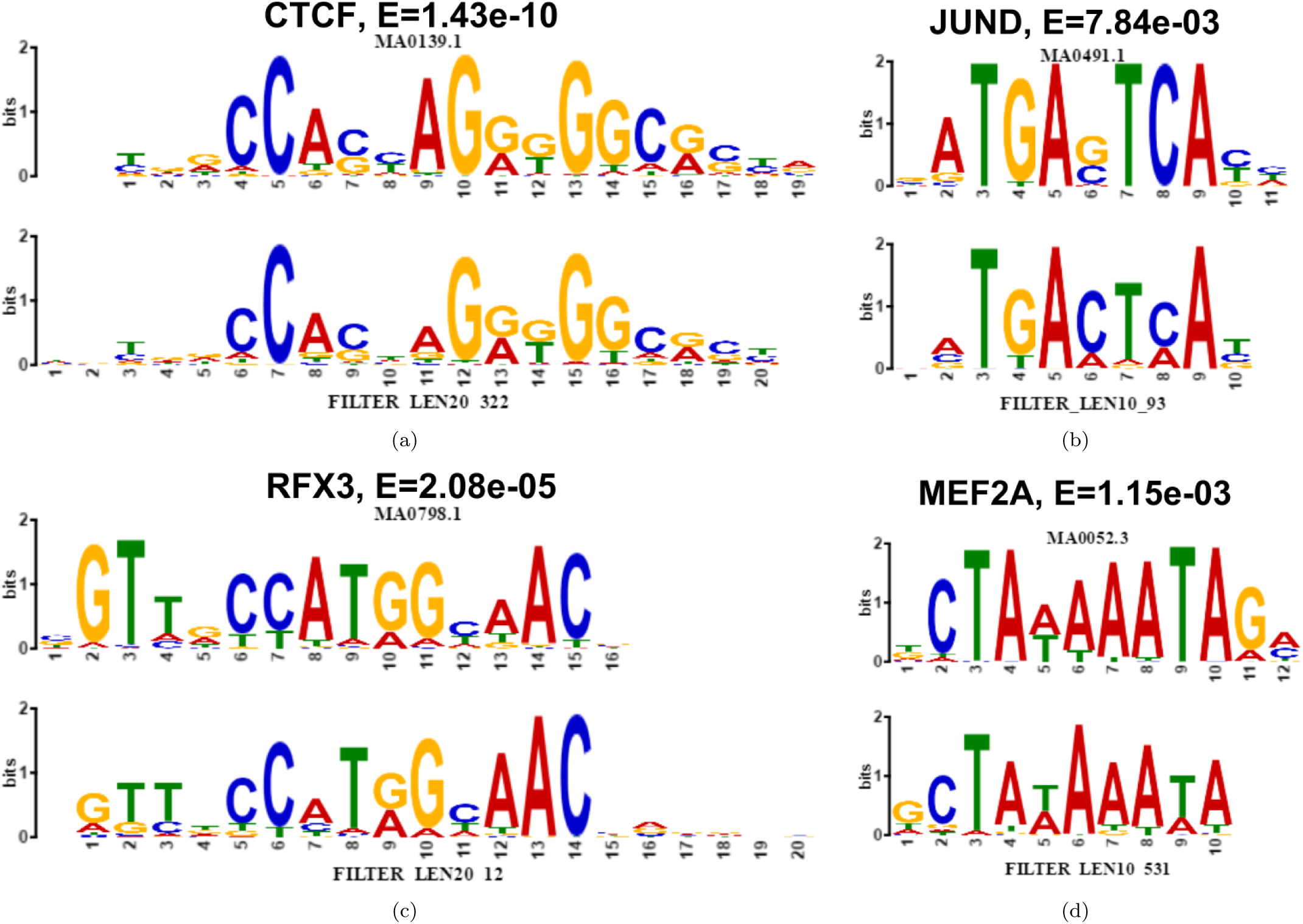
Four known motifs (top) aligned with convolution kernels (bottom). E-values of the match are displayed. (a)CTCF; (b)JUND; (c)RFX3; (d)MEF2A.

#### 3.3 Positional bias

We observed that many of the convolution kernels have display bias relative to TSS, TES and miRNA. Specifically, we downloaded RefSeq gene models[21] and obtained 4,000 bp sequences centered on the TSS or the TES of each gene. Then, we used CentriMo [22] to assess the positional bias of each kernel relative to TSS and TES. 264 and 779 kernels have significant (*E* < 1*e* – 5) positional bias relative to TSS and TES, respectively, indicating these kernels have potential roles in transcriptional control and post-transcriptional regulation. Figure 4 shows the positional distributions of the top four biased kernels relative to TSS and TES. We note that, the well known polyadenylation signal AATAAA and its reverse compliment TTTATT are among the top four positional biased kernels relative to TES. Previous results have also reported that motifs discovered in conserved sequences have positional bias relative to TSS and TES [23].

**Figure 4:**
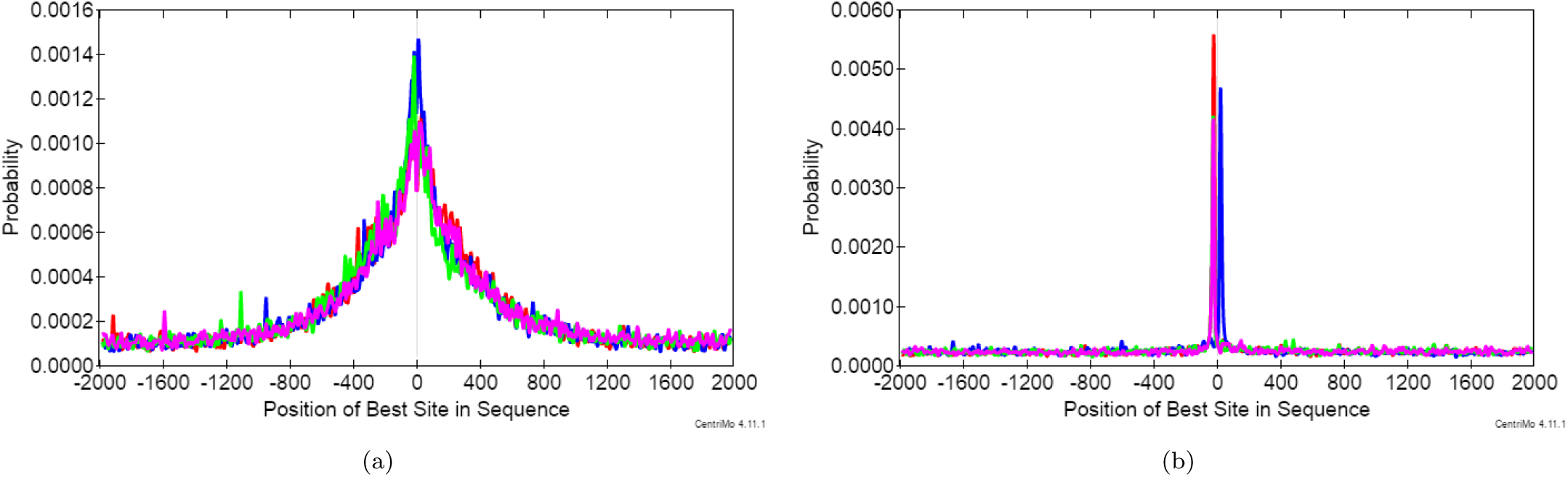
The positional distributions of the top four biased kernels relative to TSS and TES. (a)TSS; (b)TES.

Besides positional bias relative to TSS and TES, we also observed several kernels have positional bias relative to miRNA. We downloaded all the 1,881 human hairpin miRNA from miRBase [24] and used CentriMo [22] to test the positional bias of each kernel relative to miRNA. 122 kernels have significant (*E* < 1*e* – 5) positional bias relative to the first 10 positions of miRNA. Previous results have also reported that 95% of 8-mers discovered in conserved sequences match the first 10 positions of miRNA [23].

#### 3.4 Strand bias

In addition to positional bias, we also observed the convolution kernels that are close to TES have strand bias. Specifically, for each of the top 100 positional biased kernels relative to TSS and TES, we looked at the strand of the genes that the kernel is close to. Figure 5 shows the fractions of forward strand genes for each kernel. We found the fractions of forward strand genes is tightly distributed around 0.5 for kernels that positional biased to TSS, while the fractions significantly deviate from 0.5 for kernels that are positional biased to TES, suggesting those kernels also have strand bias and their RNA level regulatory effects.

**Figure 5:**
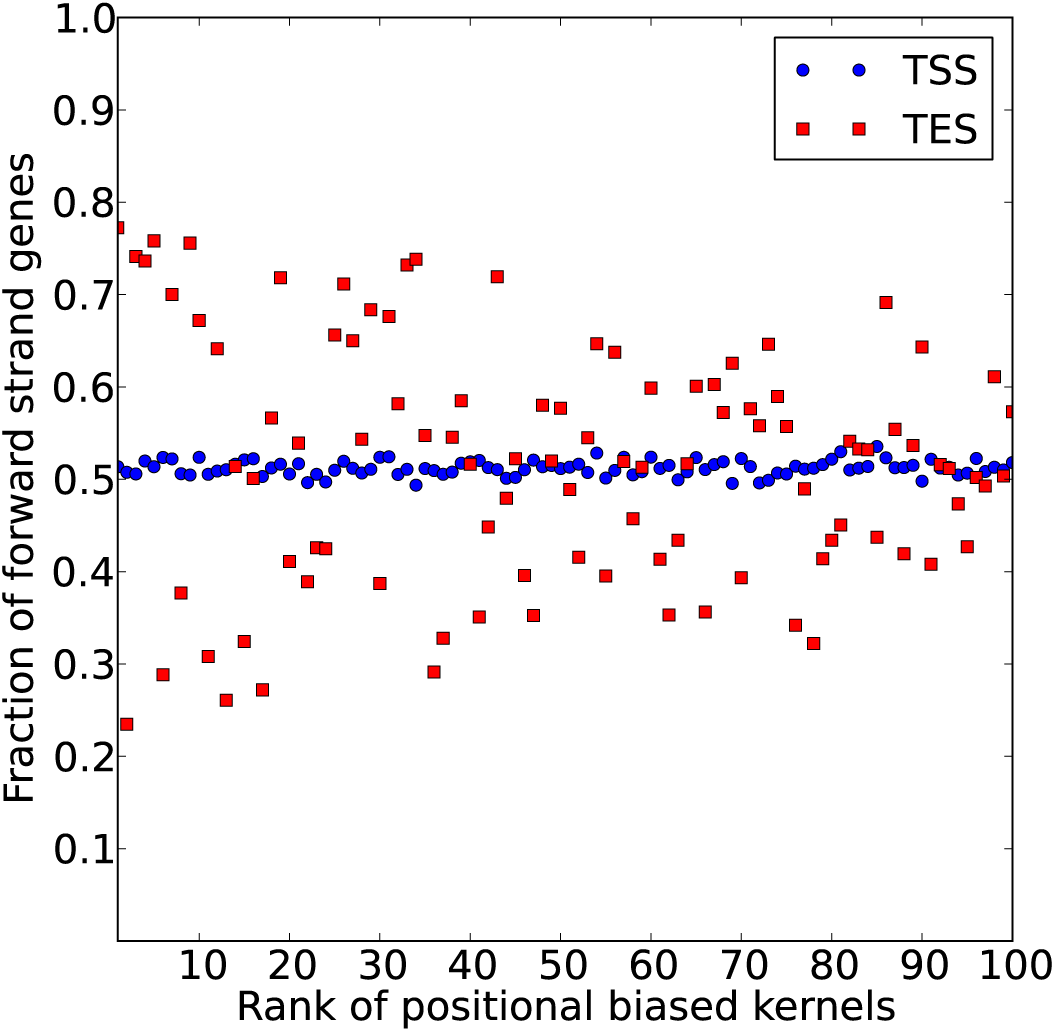
The strand bias of the top 100 positional biased kernels relative to TSS and TES. The x-axis is the rank of each kernel. The y-axis is the fraction of forward strand genes that each kernel is positional biased to.

#### 3.5 Scoring sequences at nucleotide level resolution

We adopted the method of saliency maps [25, 26] to compute the gradient of a given sequence and used it as a score to annotate each nucleotide within the sequence. Negative gradients were clipped to 0. Figure 6 shows the saliency maps of four conserved sequences. The motifs of CTCF, JUND, RFX3 and MEF2A are clearly recovered in this example, demonstrating their relevancy to conservation.

**Figure 6:**
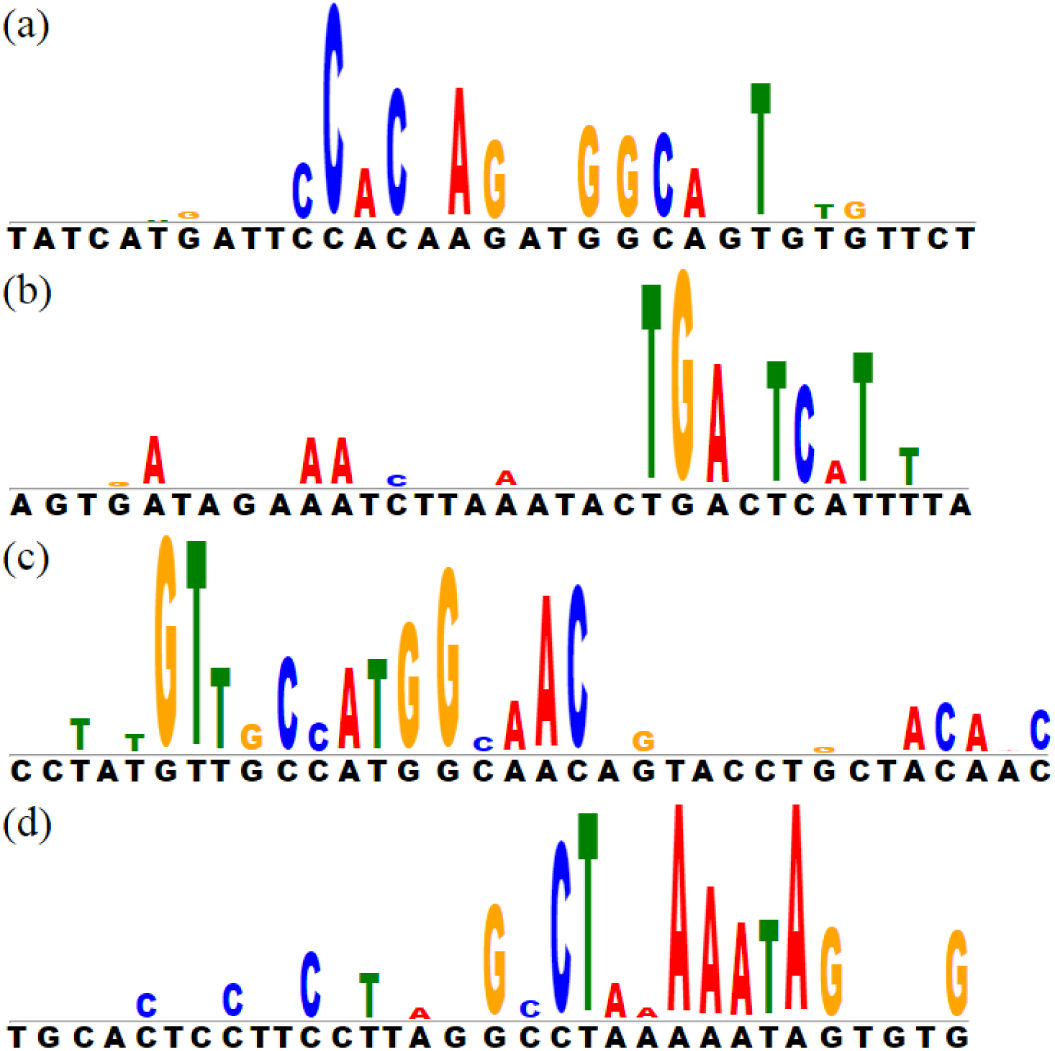
The saliency maps of four conserved sequences. The black letters below the gray line are the nucleotides of each sequence. The colored letters above the gray line are the nucleotides highlighted by their gradients, with the height proportional to the gradient. Four motifs are rediscovered in this example. (a)CTCF; (b)JUND; (c)RFX3; (d)MEF2A.

#### 3.6 Motifs summary

Finally, we clustered all 1,500 convolution kernels into 820 clusters, using RSAT motif hierarchical clustering tool [27]. The clustering results suggest that DeepCons learned a large variety of informative motifs (Figure 7). Lots of the motifs have been characterized in previous sections, while most of other motifs do not correspond to any known functional element, suggesting that they might represent unknown categories of functional elements. The complete RSAT clustering results and the 1,500 kernels in the format of MEME [13] are publicly available online at https://github.com/uci-cbcl/DeepCons.

**Figure 7:**
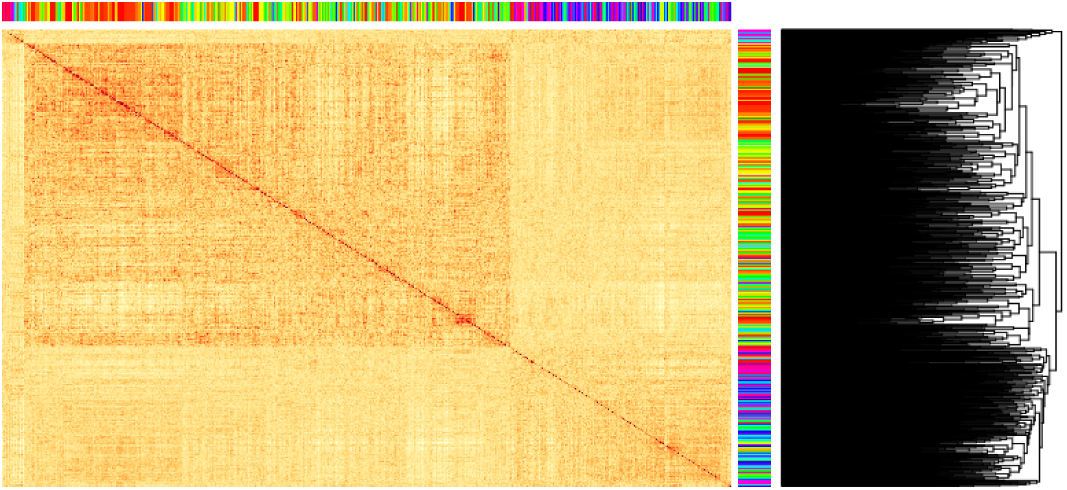
The hierarchical clustering heatmap of all the 1,500 kernels using RSAT motif clustering tool [27]

### 4 Discussion

Comparative genomics is an powerful tool in finding functional elements across the human genome. However, our understanding of the functional roles of these conserved sequences remains incomplete, especially in noncoding regions. Here we present a deep learning approach, DeepCons, to understand sequence conservation by training a convolutional neural network to classify conserved and non-conserved sequences. The learned convolution kernels of DeepCons captured rich information with respect to sequence conservation that 1) some of them match to known motifs that are known to be widely distributed within conserved noncoding elements, 2) lots of them have positional bias relative to both transcription start sites (TSS), transcription end sites (TES) and miRNA, indicating their potential roles in transcriptional control and post-transcriptional regulation regulation, and 3) they also have strand bias relative to TES, suggesting their RNA level regulatory effects. Given that the kernels alone can capture known sequence biases in some annotated genic elements, it may be interesting to understand and visualize the subsequent layers. One promising solution is to apply a dimension reduction on the activations from the dense layer and observe whether other non-coding elements such as long non-coding RNAs, TSSs, and TESs cluster together. Other clusters may correspond to functional elements that have yet to be properly characterized. We finally demonstrate that DeepCons could be used to score sequence conservation at nucleotide level resolution. We rediscovered known motifs, such as CTCF, JUND, RFX3 and MEF2A, within a given sequence by highlighting each nucleotide regarding their scores. In summary, we have developed a new deep learning framework for studying conserved sequences using convolutional neural networks. We have made all the kernels publicly available online at https://github.com/uci-cbcl/DeepCons as motifs, and we hope researchers may discover new biology by studying these motifs.

Convolutional neural networks are very effective at finding local sequence patterns through its kernels, but the kernels will typically fail to find long range sequence patterns that correspond to complex regulatory mechanisms. The size of the pattern mostly depends on the length of the convolution kernel, which typically ranges from a few bases to less than one hundred bases. Using multiple convolutional layers may help to capture broader ranges of sequence patterns, but interpreting kernels at top layers that are not directly connected to the input sequences remains difficult. Long short term memory (LSTM) networks [28], on the other hand, are Specifically designed to capture long term sequential patterns, and have been widely applied to analysis natural languages [29]. However, LSTM is also very inefficient to train since its backpropagation step is equivalent to passing the error through dozens, even hundreds, of layers. We applied LSTM to classify conserved and non-conserved sequences, but due to the large training set the algorithm took prohibitively long time to just finish even one epoch. Next, we plan to investigate multi-GPU training schemes that are now supported by TensorFlow [30], and hopefully this solution will speed up training LSTM to within an acceptable time range. Interpreting LSTM trained on sequence data also requires novel thinking. Visualizing the memory cell activities [31] may shed some lights on revealing long term sequence patterns.

## 5 Supplementary

### 5.1 Parameter configurations

DeepCons is composed of:

1. Convolutional layer of 1,000 kernels of length 10 and 500 kernels of length 20 with rectified linear activation.
2. Global max-pooling layer, with window size and stride size equals to the sequence length.
3. Fully connected dense layer of 1,500 hidden units with rectified linear activation.
4. Sigmoid output layer with cross-entropy loss.

25% dropout [32] was applied to the convolutional layer and 50% dropout was applied to the fully connected dense layer. Adagrad [15] combined with mini-batch stochastic gradient descent of batch size 128 was used to perform parameters update. Model training was scheduled for 100 epochs but may early stop if the validation loss did not decrease for 10 epochs.

### 5.2 Data preprocessing

To build the dataset of conserved sequences, we downloaded the 46-way phastCons conserved elements [1] under mammal category from UCSC genome browser [19] based on hg19. The original conserved elements were further flanked 5 bp both upstream and downstream, and overlapping elements after flanking were merged into one element. We excluded conserved sequences that overlap with repetitive sequences (http://www.repeatmasker.org/) or coding exons. We then filtered away conserved sequences that were either shorter than 30 bp or longer than 1,000 bp for training the model, leaving 887,577 sequences in the end. 75% of the nucleotides were preserved after the length filtering. To build the dataset of non-conserved sequences, we randomly shuffled the 887,577 conserved sequences on hg19, excluding repetitive sequences, coding exons and conserved sequences themselves. We then obtained the reverse complement of each sequence. Both the original sequence and the reverse complement were padded with the letter “N” such that all the sequences have the same length of 1,000 bp, and were finally concatenated with a gap of 20 bp “N”. Raw sequences in text format were converted into Numpy format with one-hot encoding, which are the input format for model training. We randomly set aside ∼80% for training (1,415,154 sequences), ∼10% for validation (180,000 sequences) and ∼10% for testing (180,000 sequences).

